# ADAM17 inhibition prevents neutrophilia and lung injury in a mouse model of Covid-19

**DOI:** 10.1101/2021.04.10.439288

**Authors:** Nathaniel L. Lartey, Salvador Valle-Reyes, Hilda Vargas-Robles, Karina E. Jiménez-Camacho, Idaira M. Guerrero-Fonseca, Ramón Castellanos-Martínez, Armando Montoya-García, Julio García-Cordero, Leticia Cedillo-Barrón, Porfirio Nava, Jessica G. Filisola-Villaseñor, Daniela Roa-Velázquez, Dan I Zavala-Vargas, Edgar Morales-Ríos, Citlaltepetl Salinas-Lara, Eduardo Vadillo, Michael Schnoor

## Abstract

Severe coronavirus disease 2019 (Covid-19) is characterized by lung injury, cytokine storm and increased neutrophil-to-lymphocyte ratio (NLR). Current therapies focus on reducing viral replication and inflammatory responses, but no specific treatment exists to prevent the development of severe Covid-19 in infected individuals. Angiotensin-converting enzyme-2 ACE-2) is the receptor for SARS-CoV-2, the virus causing Covid-19, but it is also critical for maintaining the correct functionality of lung epithelium and endothelium. Coronaviruses induce activation of a disintegrin and metalloprotease 17 (ADAM17) and shedding of ACE-2 from the cell surface resulting in exacerbated inflammatory responses. Thus, we hypothesized that ADAM17 inhibition ameliorates Covid-19-related lung inflammation. We employed a pre-clinical mouse model using intra-tracheal instillation of a combination of polyinosinic:polycytidylic acid (poly-I:C) and the receptor-binding domain of the SARS-CoV-2 spike protein (RBD-S) to mimic lung damage associated with Covid-19. Histological analysis of inflamed mice confirmed the expected signs of lung injury including edema, fibrosis, vascular congestion and leukocyte infiltration. Moreover, inflamed mice also showed an increased NLR as observed in critically ill Covid-19 patients. Administration of the ADAM17 inhibitors apratastat and TMI-1 significantly improved lung histology and prevented leukocyte infiltration. Reduced leukocyte recruitment could be explained by reduced production of pro-inflammatory cytokines and lower levels of the endothelial adhesion molecules ICAM-1 and VCAM-1. Additionally, the NLR was significantly reduced by ADAM17 inhibition. Thus, we propose inhibition of ADAM17 as a novel promising treatment strategy in SARS-CoV-2-infected individuals to prevent the progression towards severe Covid-19.

## Introduction

Severe acute respiratory syndrome coronavirus 2 (SARS-CoV-2) is the infectious agent that causes coronavirus disease 2019 (Covid-19), which can cause lung failure in susceptible individuals (1). This virus has infected 129,750,743 people and caused 2,830,016 deaths worldwide (Source: https://coronavirus.jhu.edu/map.html; 04/02/2021). SARS and severe Covid-19 are characterized by pulmonary edema, fibrosis, activation of the coagulation cascade, hypoxia, and hypercapnia (2). Moreover, severe Covid-19 is characterized by neutrophilia and lymphopenia and thus increased neutrophil-to-lymphocyte ratio (NLR) (3, 4). Excessive neutrophil presence and recruitment to the lung and transmigration into the alveolar luminal space contribute to lung injury as neutrophils release proteases, reactive oxygen species and neutrophil extracellular traps (NETs), which are crucial to combat pathogens, but in large amounts are harmful to the host tissue (5). Survival depends on the severity of lung injury, the extension of damage to other organs, comorbidities of the infected patient, and the quality of medical care (6).

SARS-CoV-2, via its surface protein Spike, uses angiotensin-converting enzyme-2 (ACE2) for host entry via airway epithelial cells (7). ACE2 is an enzyme highly expressed on the apical side of polarized alveolar epithelial cells and lung endothelial cells that cleaves the peptide hormone angiotensin-I into Ang(1-9) and Ang(1-7), peptides known to protect against epithelial and endothelial hyperpermeability and excessive inflammation (8). Genetic deletion of ACE2 worsens lung inflammation, while Ang(1-7) administration improve it (9). Coronavirus infection causes downregulation of ACE2 by activating enzymes known to cleave ACE2 (10) including a disintegrase and metalloprotease 17 (ADAM17) (11, 12). ADAM17 sheds the extracellular domain of ACE2 from the surface of lung epithelial cells thus reducing the protective ACE-2-dependent signaling leading to a detrimental feedback loop of exacerbated lung inflammation. Moreover, ADAM17 produces the active form of the pro-inflammatory cytokine tumor necrosis factor-α (TNF-α) and is critical for proinflammatory cytokine secretion thus contributing to the cytokine storm, another feature of severe Covid-19 that triggers excessive neutrophil recruitment (13-15). Although it is well-known that ADAM17 gets activated after SARS-CoV infection (12), cleaves ACE-2, and is therefore likely involved in Covid-19 pathogenesis (16), the role of ADAM17 in the development of severe Covid-19 has not been experimentally analyzed.

Pharmacological therapy of critically ill patients relies on inhibiting viral replication and inflammatory responses (17). Although the development of vaccines has been astonishingly fast and much progress has been made with vaccinations worldwide, specific treatments to prevent the development of severe Covid-19 in infected individuals are still desperately needed for those that do not wish to or cannot be vaccinated, or those that get infected despite being vaccinated.

We hypothesized that pharmacological inhibition of ADAM17 protects from acute lung injury (ALI) by preventing excessive cytokine production and uncontrolled neutrophil recruitment. We show that intraperitoneal or intranasal administration of the ADAM17 inhibitors apratastat and TMI-1 significantly reduced proinflammatory cytokine production and leukocyte recruitment to the lungs that was accompanied by improved lung histology. Our data suggest that inhibition of ADAM17 is a viable strategy to prevent severe Covid-19 cases and thus reducing the requirement for critical care and Covid-19 death toll.

## Methods

### Experimental Animals

Pathogen-free 8-12 weeks old C57BL/6 male mice were obtained from the animal facility in CINVESTAV. All protocols have been approved by the institutional animal care and use committee of CINVESTAV.

### Production, purification and refolding of the RBD-S protein

The recombinant receptor-binding domain (RBD) of the SARS-CoV-2 Spike protein (RBD-S) with amino and carboxy deletions (RDB-NTCT) was purified from *E. coli* SoluBL21 (DE3) (Genlantis) cells transformed with pRSET-RDB-NTCT. Briefly, cells were grown in 20 ml LB broth supplemented with ampicillin (100 µg/ml) at 37 °C for 12 h at 150 rpm. This 20 ml of culture was added into 500 ml of 2xYT broth (tryptone16 g/L, yeast extract 10 g/L and sodium chloride 5 g/L) and ampicillin (100 μg/ml), incubated at 37 °C under constant shaking at 200 rpm until reaching an OD_600_ of 0.8. Expression of the protein was induced by adding extra 500 ml of 2xYT broth with 2 mM isopropyl β-d-galactopyranoside and 2 mM MgSO_4_ (to reach a final volume of 1 L of culture and 1 mM IPTG and 1 mM MgSO_4_). The bacteria were incubated at 16 °C for 16 h and harvested by centrifugation at 2500 x g for 30 min at 4 °C. The bacterial pellet was resuspended in 20 ml of lysis buffer (20 mM Tris-HCl pH 8.0, 100 mM NaCl, 1 mM PMFS, 1 mM benzamidine and 1 mM DTT) and sonicated in an ice bath using an Ultrasonic Homogenizer (500-Watt Cold Palmer, USA) for 4 min. The cell lysate was centrifuged 24000 x g for 20 min at 4 °C. The purification of the RBD-NTCT domain from inclusion bodies was performed as described previously (18) with the following modifications. The insoluble fraction was homogenized using an ULTRA-TURRAX T18 homogenizer (IKA, Germany). Inclusion bodies were solubilized in buffer A (20 mM Tris-HCl pH 8.0, 500 mM NaCl, 5 mM imidazole, 1 mM DTT and 8 M Urea) and incubated for 16 h at 20 °C with constant agitation at 150 rpm. The solubilized inclusion bodies (supernatant) were recovered by ultracentrifugation at 118,000 x g for 30 min at 4 °C. The supernatant was filtered through a 0.45 μm membrane and applied onto a HisTrap Ni column (5 ml, Cytiva) previously equilibrated with buffer A with a FPLC (ÄKTApure 25, Cytiva). After loading the sample, the column was washed with the same buffer to eliminate unbound proteins and subsequently the RBD-NTCT domain was eluted through a linear gradient with buffer B (20 mM Tris -HCl pH 8.0, 500 mM NaCl, 500 mM imidazole, 1 mM DTT and 8 M urea). Refolding of the RBD-NTCT domain was performed by size exclusion chromatography with a Superdex 200 Increase 10/300 GL column (Cytiva). The column was equilibrated with refolding buffer (20 mM Tris-HCl pH 8.0, 100 mM NaCl, 5 % glycerol, 0.4 oxidized glutathione, 0.2 mM reduced glutathione, 1 mM PMSF and 100 mM arginine). For each refolding run, we applied 1 ml of sample from the previous purification step onto the column followed by a linear gradient from 8 M urea to refolding buffer. The protein was eluted with the refolding buffer. The buffer of the refolded protein was replaced with PBS using a Hi Trap desalting column (5 ml, Cytiva) for subsequent experiments.

### Preclinical model of lung inflammation and ADAM17 inhibitor administration

This Covid-19-related pre-clinical model of lung inflammation is based on a recently published protocol with the exception that we here used the recombinant receptor-binding (RBD) domain of the SARS-CoV-2 Spike protein instead of the entire extracellular domain (19). Mice were anesthetized by intraperitoneal injection of 200 mg/kg ketamine (Anesket, PISA, Mexico-City) and 10 mg/kg xylazine (Procin, PISA, Mexico-City). The trachea was surgically exposed and a sterile 31G needle was carefully inserted. A combination of 10 mg/kg poly-I:C (Invivogen, San Diego, CA) and 15 μg of the recombinant RBD of SARS-CoV-2 Spike protein (RBD-S) in 60 μL endotoxin-free physiological saline was instilled followed by 100 μL of air to fill the lungs. Sham-operated animals received physiological saline alone as control. The wounds were sutured, and the mice left to recover in cages. The ADAM17 inhibitors apratastat and TMI-1 (Sigma Aldrich-Merck, Toluca, Mexico) were administered either intraperitoneally or intranasally at 10 mg/kg (20, 21). Apratastat was administered twice (4h and 16 h after surgery) and TMI-1 was administered once (4 h after surgery). DMSO was administered as vehicle control. Twenty-four h after induction of lung inflammation, blood samples (cardiac puncture), lung tissue and bronchoalveolar lavage fluid (BALF) were obtained for analysis of inflammation as described below.

### Histology

After obtaining blood by cardiac puncture, 20 ml of PBS was transcardially perfused until the lungs turned white. The right lung was extracted and fixed in 4% formalin. Fixed lung specimens were dehydrated, embedded in paraffin. 4 μm cross-sections were stained with haematoxylin and eosin according to standard protocols. A pathologist analyzed the samples for the degree of inflammation in a blinded fashion and determined the histopathological scores as previously published (22).

### Bronchoalveolar lavage

24 h after induction of lung inflammation, mice were euthanized by anesthesia overdose and bronchoalveolar lavage was performed as described before (23). Briefly, the rib cage was removed, and a plastic cannula was inserted into the trachea and lungs were flushed three times with 0.8 ml PBS. This procedure was repeated four times. Pooled BALF was analyzed by flow cytometry as described below.

### Lung digestion

Left lungs were put in 2 ml microtubes with 1 ml prewarmed PBS containing 200 units of collagenase (Sigma, C9891) and 0.9 mM of calcium and magnesium. Tissue was cut into pieces using sharp scissors. A small hole was made in the cap of the microtubes to prevent hypoxia and the lung suspension was incubated for 20 min at 37 °C in a water bath. Samples were vigorously vortexed and then resuspended using a pipette tip. Lung tissue suspensions were then incubated for 20 min more at 37°C. The cell suspension was then filtered through a 40 μm cell strainer (Corning) and washed with 5 ml of PBS. Cells were then centrifuged at 180 x g for 5 minutes and resuspended in 1 ml PBS for staining and analysis by flow cytometry.

### Flow Cytometry

Cells from blood, BALF and lung suspension were blocked for 15 min with anti-mouse TruStain FcX (Biolegend, Mexico-City) and stained with anti-mouse CD45 Pacific blue (total leukocytes), anti-mouse Ly6G APC-Cy7 (neutrophils), anti-mouse F4/80 FITC (macrophages), anti-mouse CD3 PE (lymphocytes) and anti-mouse PECAM-1 and ICAM-1 (endothelial cells) for 15 min. Data were acquired using a FASC Canto II and analyzed using FlowJo V10 software.

### RNA isolation, cDNA synthesis and qRT-PCR

Total RNA was isolated and purified from lung tissues using Trizol Reagent (Invitrogen, CA). cDNA was obtained by reverse-transcribing the purified RNA using the first stand cDNA synthesis kit (Thermofisher, Massachusetts). The following primer pairs were used: *β-actin*: forw 5′-TATCCACCTTCCAGCAGATGT-3′; rev. 3′-AGCTCAGTAACAGTCCGCCTA-5′. *TNF-α*: forw. 5′-ACGGCATGGATCTCAAAGAC-3′; rev. 3′-AGATAGCAAATCGGCTGACG-5′. *IL-1β*: forw. 5′-GCAACTGTTCCTGAACTCAACT-3′; rev. 3′-TCTTTTGGGGTCCGTCAACT-5′. *IL-6*: forw. 5′-CCTTCCTACCCCAATTTCCAA-3′; rev. 3′-AGATGAATTGGATGGTCTTGGTC-5′. *IL-10*: forw. 5′-ACTGCACCCACTTCCCAGT-3′; rev. 3′-TGTCCAGCTGGTCCTTTGTT-5′. *ICAM-1*: forw. 5′-CAATTTCTCATGCCGCACAG-3′; rev. 3′-AGCTGGAAGATCGAAAGTCCG-5′. *VCAM-1*: forw. 5′-TGAACCCAAACAGAGGCAGAGT-3′; rev. 3′-GGTATCCCATCACTTGAGCAGG-5′. PCR was performed using Power SYBR Green PCR master mix (2X) in a final volume of 10 μL. The reaction mixture contained 5.0 μL of the master mix, 25 ng cDNA, and 0.15 μg of each primer. PCR reactions were run in a StepOneTM Real-Time PCR System. PCR conditions were as follows: activation for 10 min at 95 °C; 40 cycles of (denaturation at 95 °C for 15 s and data acquisition during annealing and extension at 60 °C for 60 s). The melting curve was generated afterward by heating at a rate of 0.6 °C/s from 60 °C to 95 °C. Using β-actin as endogenous control, expression was quantified using the 2^-ΔΔCt^ method.

### ELISA

The levels of serum TNF-α were determined using the LEGEND MAX Mouse TNF-α ELISA kit (Biolegend, CA) according to the manufacturer′s instructions.

### Statistical Analysis

Data were analyzed using GraphPad Prism Version 6.0. One-way analysis of variance (ANOVA) followed by Dunnett post-hoc was used for multi-group comparisons. p-values < 0.05 were considered statistically significant.

## Results

### Preclinical model of Covid19-related lung inflammation

To test whether ADAM17 would have a beneficial effect on Covid-19-related lung inflammation, we employed a pre-clinical mouse model of ALI, in which we instilled into mouse lungs a combination of the Toll-like receptor-3 (TLR-3) ligand poly-I:C to mimic viral infection and the recombinant RBD-S to mimic SARS-CoV-2 infection. This pre-clinical Covid-19 model is based on a recently published protocol (19, 22), with the simplification that we used the Spike RBD-domain instead of the whole extracellular domain. The ADAM17 inhibitors apratastat and TMI-1 were administered either i.p. or i.n. and 24 h after induction of lung inflammation, lung and blood were harvested and bronchoalveolar lavage performed. The experimental setup is illustrated in figure 1. This model closely resembled the Covid-19 histopathology observed in severe Covid-19 patients (24, 25), including edema formation, fibrosis, and leukocyte infiltration (Fig. 2).

**Figure 1:**
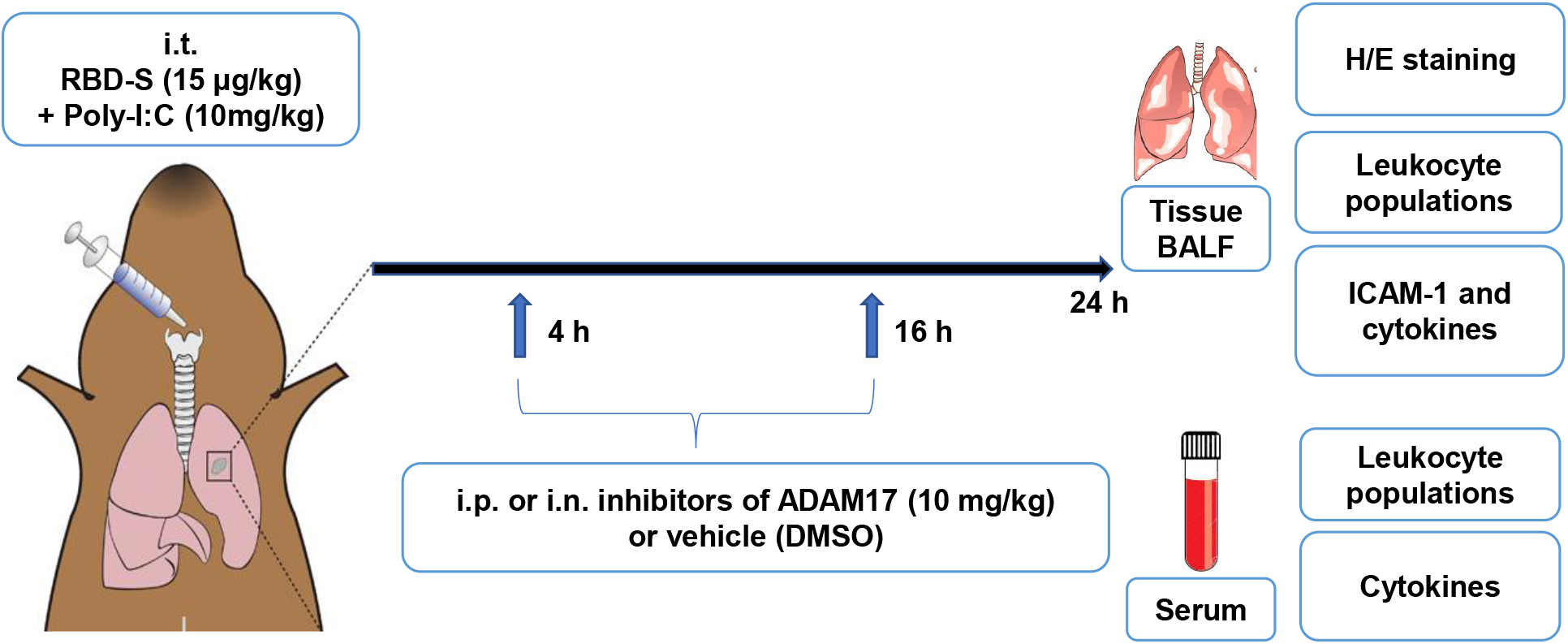
Cartoon depicting the preclinical mouse model of Covid19-related lung inflammation. The model consists of intratraqueal (i.t.) instillation of poly I:C and RBD-S, intraperitoneal (i.p.) or intranasal (i.n.) treatment with apratastat, TMI-1 or vehicle (DMSO), and harvesting of samples for the indicated analyses.

**Figure 2.**
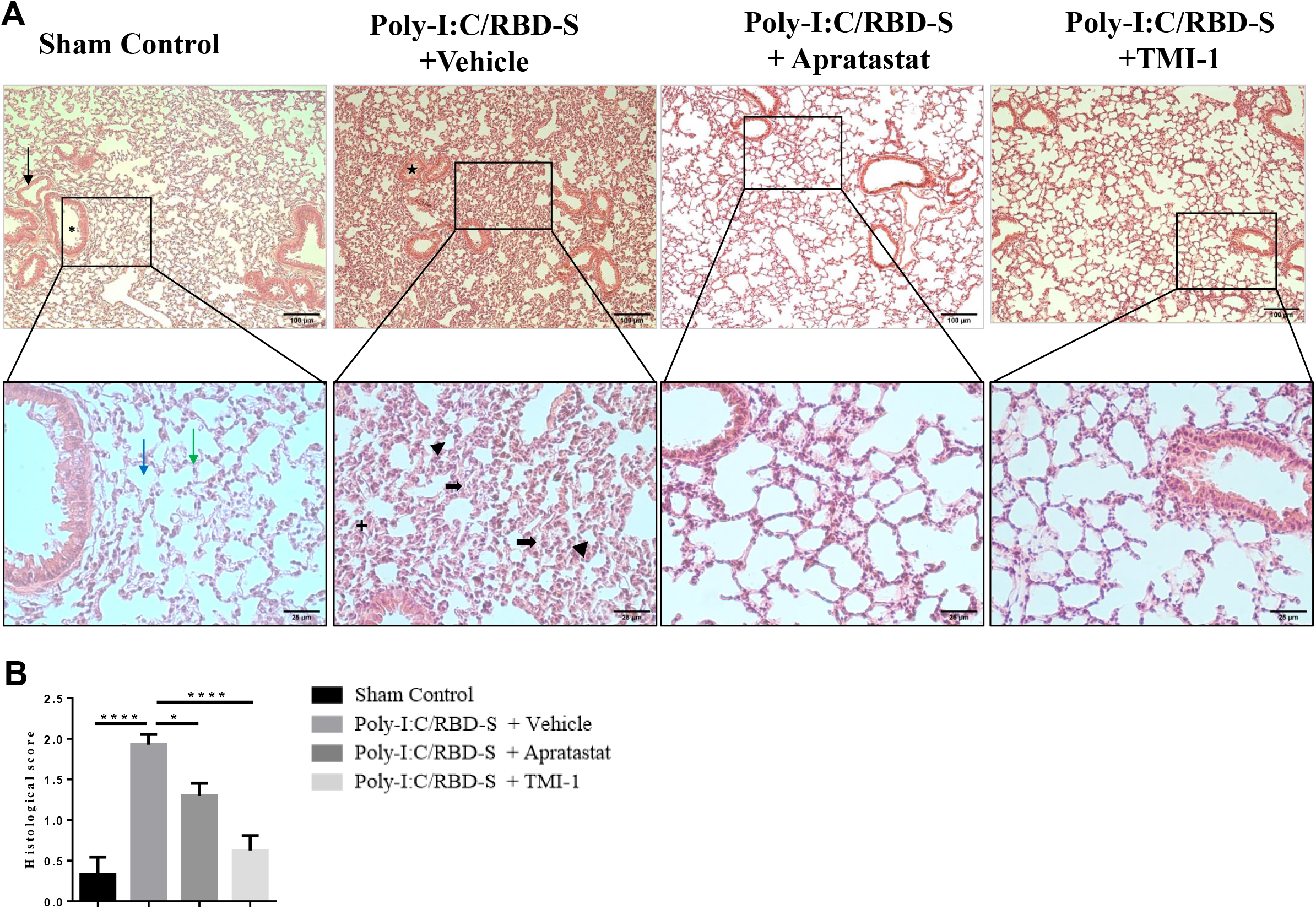
Lung injury is reduced upon ADAM17 inhibition in Poly I:C/RBD-S-induced lung inflammation. (A) Representative images of haemoxylin/eosin-stained lung tissue cross-sections showing normal architecture characterized by one cell thick alveolar wall, absence of edema, fibrosis and vascular congestion (Sham control; 10 x, upper panel, black arrow=blood vessel, asterisk=broncheole; 40x magnifications of the boxed areas, lower panel, blue arrow=alveoli, green arrow=alveolar septa). The Poly I:C/RBD-S + vehicle group shows major inflammatory changes such as alveolar wall/septa thickening (bold black arrow) and alveolar space closing and fibrosis (arrowhead), leukocyte infiltration (+) and vascular congestion (star). The lung tissues of Poly I:C/RBD-S + apratastat and + TMI groups clearly show less inflammatory damage and overall a better-preserved tissue morphology. (B) Histological scores of the lung tissues based on the extent of inflammatory changes with 0=absence, 1=low, 2=moderate, 3=high. Data are shown as mean ± SEM of images from at least 4 independent tissue preparations. *p < 0.05, ****p < 0.001.

### ADAM 17 inhibition improves lung inflammation induced by intratracheal instillation of poly I:C and RBD-S

Intratracheally instillation of the combination poly-I:C/RBD-S induced strong signs of lung inflammation and acute lung injury such as edema, fibrosis, alveolar thickening, closing of alveolar spaces and leukocyte infiltration (Fig 2A). Lung tissues of mice that received ADAM17 inhibitors showed better-preserved lung architecture and a clear amelioration of the above-mentioned clinical signs of lung injury (Fig. 2A). The protective effect seemed to be more pronounced with TMI-1 compared to apratastat. Histological scoring confirmed these impressions. Both apratastat and TMI-1 significantly reduced the histopathological scores compared to the inflamed vehicle-treated control mice, with TMI-1 showing more pronounced protection (Fig. 2B).

### Neutrophil infiltration into lungs is attenuated upon ADAM17 inhibition

Massive neutrophil infiltration is a key feature of Covid19-associated lung injury (26, 27). Because the histology indicated reduced leukocyte presence in lungs after ADAM17 inhibition, we quantified leukocyte populations by flow cytometry. As expected, the number of neutrophils was strongly increased after instillation of poly-I:C/RBD-S, and both apratastat and TMI-1 significantly reduced the number of neutrophils in the lungs (Fig. 3A). Also, the number of lung macrophages was much higher in inflamed lungs compared to controls, and again both apratastat and TMI-1 significantly reduced macrophage numbers in the lungs (Fig. 3B). By contrast, numbers of T cells were significantly reduced in inflamed lungs, but numbers remained similar after treatment with either apratastat or TMI-1 (Fig. 3C).

**Figure 3.**
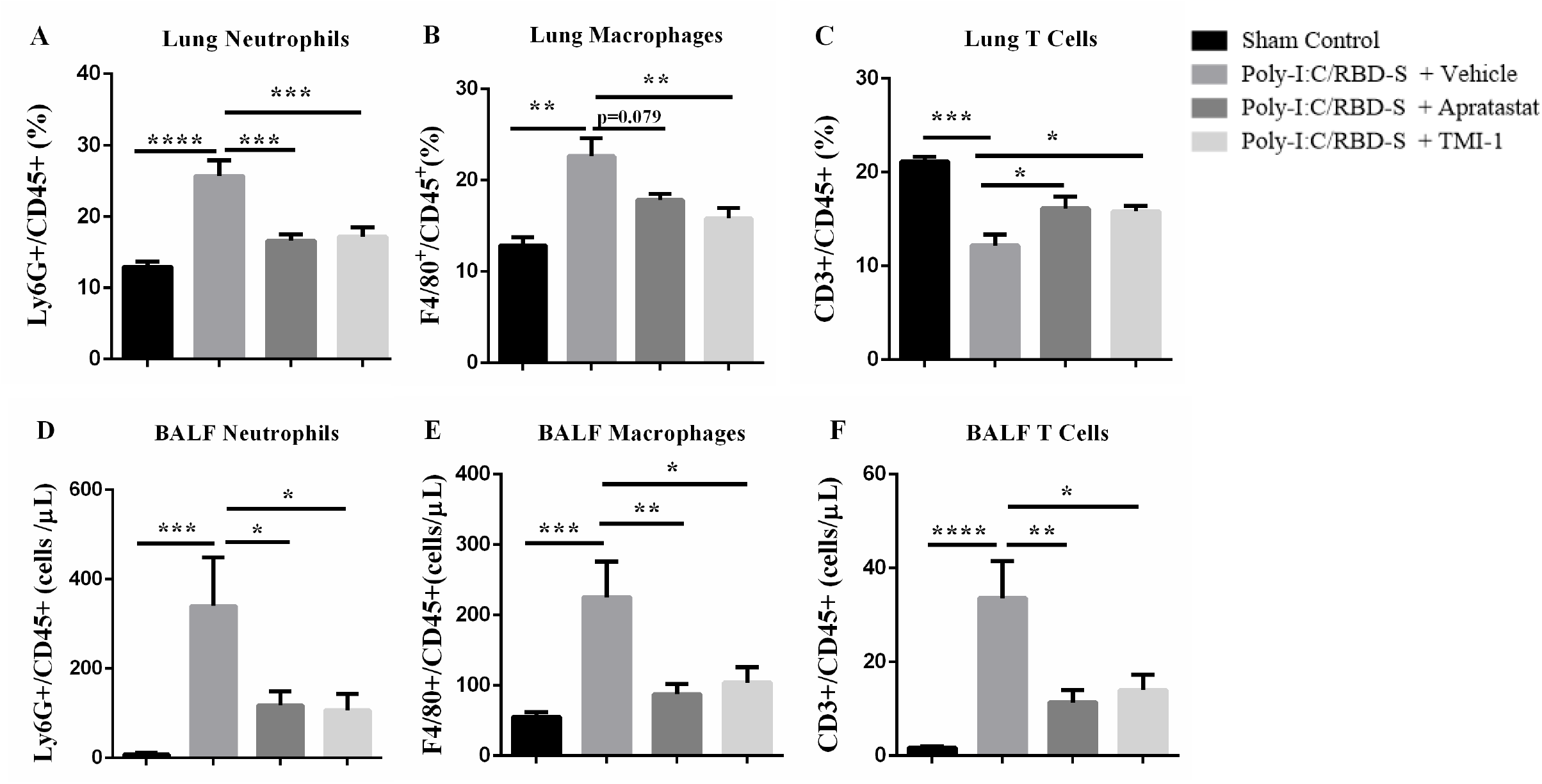
Neutrophil influx into lungs is reduced upon ADAM17 inhibition. (A) Frequency of Ly6G^+^CD45^+^ cells in the lungs of the indicated groups. (B) Frequency of F4/80^+^CD45^+^ cells in the lungs. (C) Frequency of CD3^+^CD45^+^ cells in the lungs. Cell frequencies were determined by flow cytometry 24 h after surgery. Total numbers of (D) Ly6G+ neutrophils, (E) F4/80+ macrophages, (F) CD3+ T cells from bronchoalveolar lavage fluids (BALF) 24 h after surgery. Data are shown as mean ± SEM of at least 6 mice per group. *p < 0.05, **p < 0.01, ***p < 0.001. ****p < 0.001.

Next, we analyzed leukocyte populations in the BALF, i.e. those leukocytes that transmigrated the epithelium to reach the alveolar lumen (28). We found similar results for neutrophils (Fig. 3D) and macrophages (Fig. 3E) in the BALF as in lung tissues. The strong increase in neutrophil and macrophage numbers after instillation of poly-I:C/RBD-S were significantly reduced by the ADAM17 inhibitors apratastat and TMI-1. By contrast, T cell numbers in the BALF were strongly increased in response to poly-I:C/RBD-S and were significantly reduced after ADAM17 inhibition, with both apratastat and TMI-1 showing similar effects (Fig. 3F).

### ADAM17 inhibition reverses neutrophilia and lymphopenia in poly-I:C/RBD-S-induced lung inflammation

Increased NLR is a well-established risk factor for severe Covid-19 (26). Given the changes in leukocyte numbers in the lung, we next analyzed neutrophil and T cell numbers in the blood. We found that poly-I:C/RBD-S-induced lung inflammation caused neutrophilia that was clearly ameliorated by ADAM17 inhibition (Fig. 4A). Also, we observed strong lymphopenia in response to poly-I:C/RBD-S instillation; however, T cell numbers were not changed after ADAM17 inhibition (Fig. 4B). The reduced neutrophil numbers in combination with unaltered T cell numbers showed a strong and significant reduction of the NLR (Fig. 4C), which should clearly contribute to the protective effect observed with ADAM17 inhibition.

**Figure 4.**
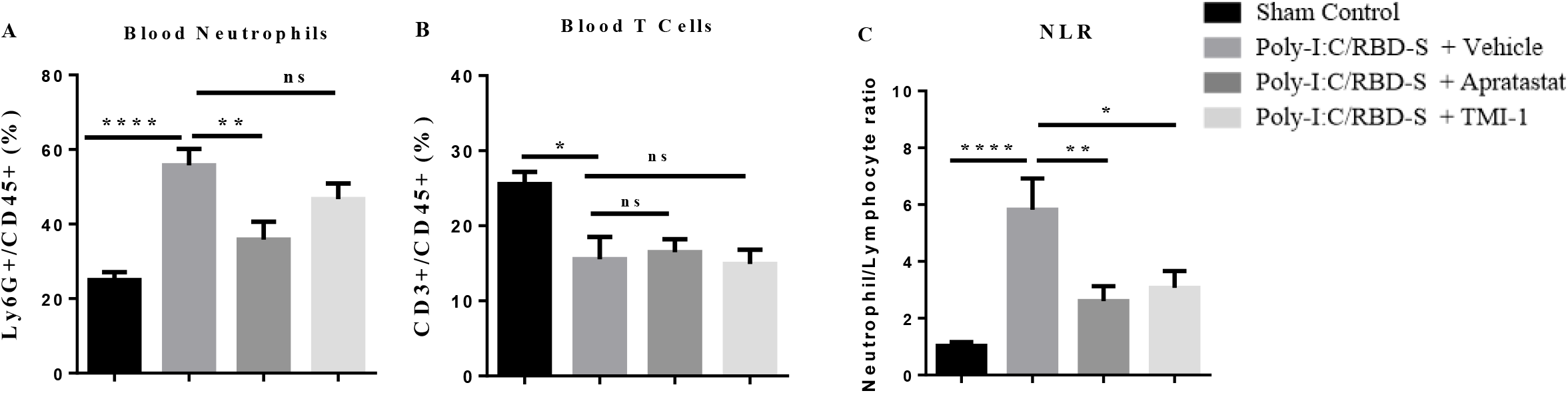
Poly-I:C/RBD-S-induced neutrophilia is prevented upon inhibition of ADAM17. (A) Frequency of Ly6G^+^ CD45^+^ neutrophils and (B) frequency of CD3+CD45+ T cells in the peripheral blood 24 h after surgery in the blood. (C) Neutrophil-lymphocyte ratio (NLR) in the peripheral blood 24 h after surgery in the blood. Data are represented as mean ± SEM of at least 6 mice per group. * p < 0.05 **p < 0.01.

### ADAM17 inhibition reduces poly-I:C/RBD-S-induced overexpression of TNF-α and endothelial adhesion molecules

Another classical feature of severe Covid-19 is the cytokine storm, which in turn leads to endothelial activation, massive leukocyte recruitment and leukocyte hyperactivation (27). ADAM17 is the enzyme responsible for releasing the active form of tumor necrosis factor-α (TNF-α) (29). Thus, we expected ADAM17 inhibition to also impact cytokine expression, endothelial activation, which in consequence is the basis for excessive neutrophil recruitment. The massive overexpression of TNF-α in response to poly-I:C/RBD-S was reverted to control levels by ADAM-17 even at the mRNA level (Fig. 5A). This was accompanied by reduced TNF-α serum protein levels (Fig. 5B) indicating that not only the local but also systemic inflammatory response is attenuated by ADAM17 inhibition.

**Figure 5.**
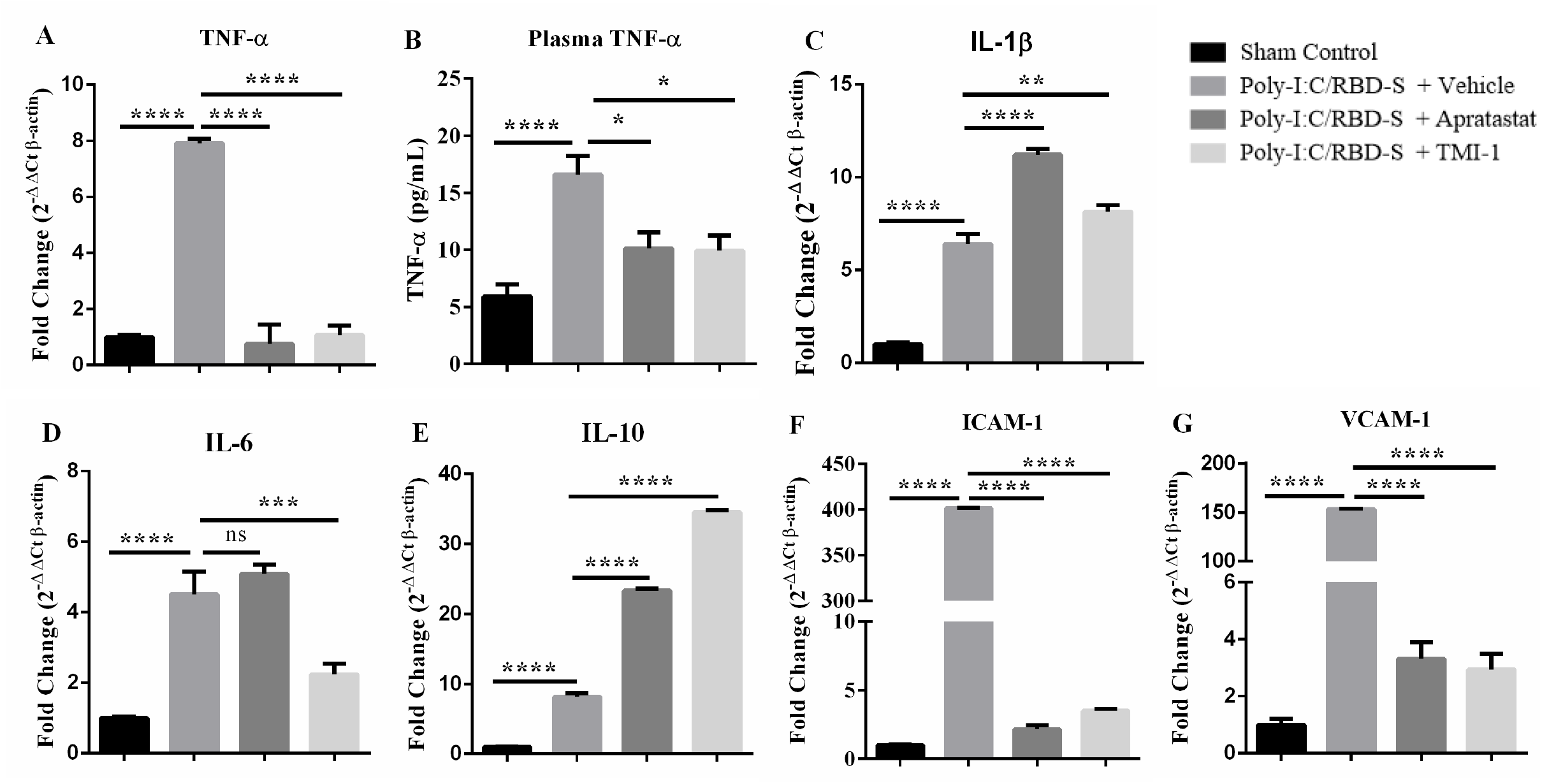
ADAM17 inhibition affects mRNA levels of cytokines and adhesion molecules. (A) Apratastat and TMI-1 ameliorate the Poly-I:C/RBD-S-induced increase in (A) TNF-α mRNA levels, and (B) TNF-α serum protein levels. (C) ADAM 17 inhibitors induce an increase in IL-1β mRNA. (D) IL-6 mRNA levels are significantly reduced only after TMI-1 treatment. (E) In Apratastat and TMI-1 further increase mRNA levels of IL-10 compared to vehicle-treated inflamed mice. The Poly-I:C/RBD-S-induced increase in mRNA levels of (F) ICAM-1 and (G) VCAM-1 are significantly reduced by treatment with both apratastat and TMI-1. Gene expression was analyzed using the quantitative real-time RT-PCR using β-actin as housekeeping gene. Data are represented as mean ± SEM of at least 4 independent cDNA preparations per group. *p < 0.05, **p < 0.01, *** p < 0.001 and **** p < 0.0001.

However, ADAM17 inhibitors further increased poly-I:C/RBD-S-induced mRNA levels of IL1-β (Fig. 5C); and only TMI-1 was able to reduce the levels of IL6 (Fig. 5D). Of note, mRNA levels of the anti-inflammatory cytokine IL10, which is known to increase during inflammation as a protective countermeasure, were significantly higher after ADAM17 inhibition (Fig. 5E), suggesting that this is another protective mechanism in the context of lung inflammation.

Pro-inflammatory cytokines, especially TNF-α, are known to activate endothelial cells that respond by expressing adhesion molecules for leukocytes to enable their extravasation (30). Accordingly, poly I:C/RBD-S induced strong expression of ICAM-1 (Fig. 5F) and VCAM-1 (Fig. 5G) in lungs; and both apratastat and TMI-1 reversed this overexpression back to control levels. This reduction in endothelial adhesion molecules likely explains the reduced leukocyte recruitment to the lungs after ADAM17 inhibition.

### Intranasal administration of ADAM17 inhibitors also prevents excessive leukocyte recruitment to the lungs in response to poly I:C/RBD-S

All data described above were obtained after intraperitoneal administration of the ADAM17 inhibitors. As this is not a desirable route of applying drugs in human patients, we wanted to know whether local application of apratastat via the intranasal route would show similar effects. Importantly, we indeed observed similar results with respect to the leukocyte populations in lung tissue and BALF. Lung neutrophils were reduced almost to control levels with ADAM17 inhibition mice (Fig 6A); lung macrophages were significantly reduced although not to control levels (Fig. 6B); and lung T cell numbers during poly-I:C/RBD-S-induced lung inflammation were increased by apratastat (Fig. 6C). Total cell numbers (Fig. 6D) and total leukocyte numbers (Fig. 6E) in the BALF were significantly reduced by apratastat during poly-I:C/RBD-S-induced lung inflammation. Analyzing the leukocyte populations in the BALF separately, we found that intranasal administration of apratastat, like intraperitoneal administration, significantly reduced the numbers of neutrophils (Fig. 6F), macrophages (Fig. 6F), and T cells (Fig. 6F).

**Figure 6.**
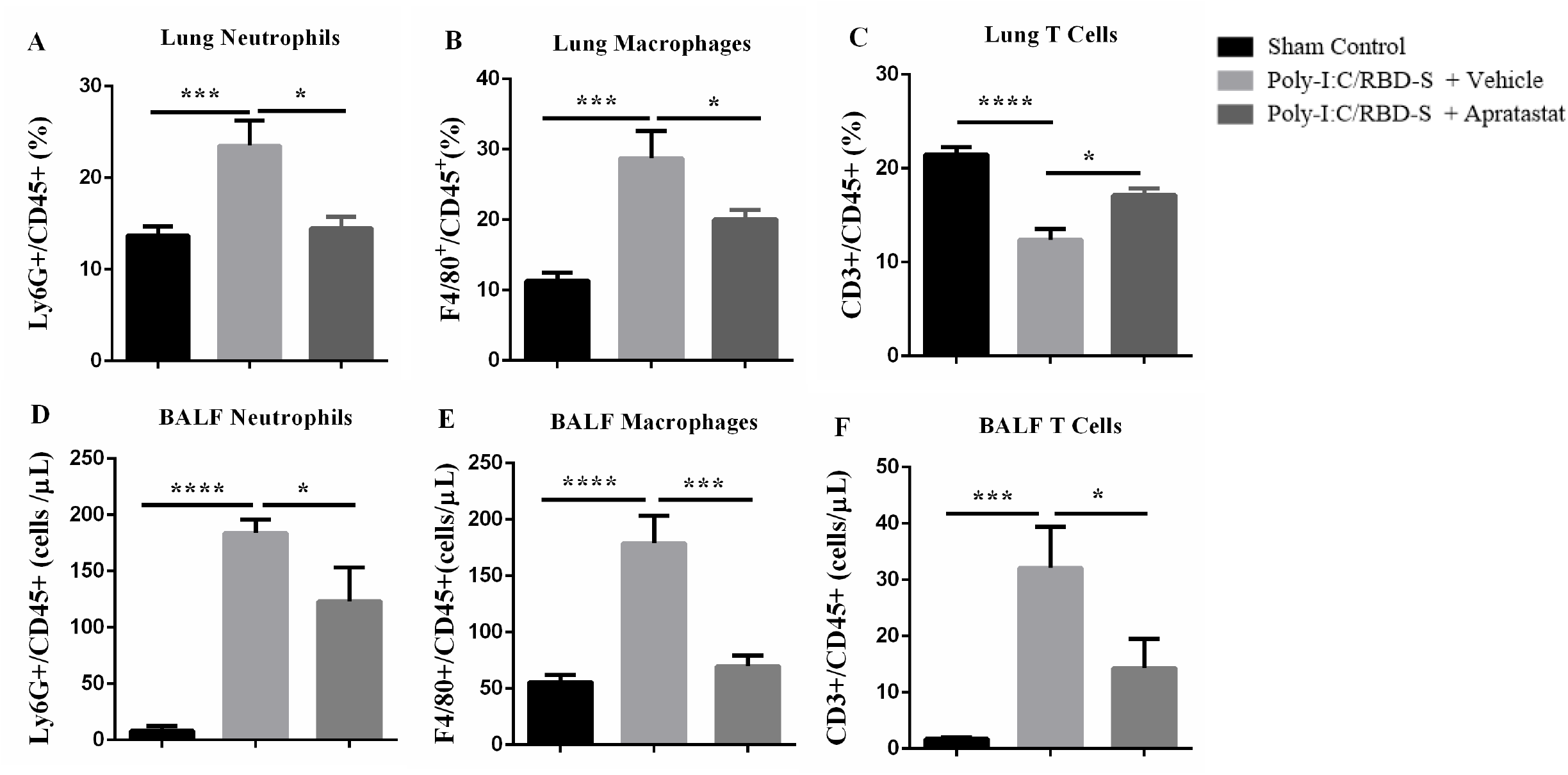
Intranasal administration of apratastat reduces neutrophil and macrophage presence in poly-I:C/RBD-S-inflamed lungs. (A) Percentage of Ly6G^+^/CD45^+^ neutrophils; (B) F4/80^+^/ CD45^+^ macrophages, and (C) CD3^+^/CD45^+^ T cells in lung tissue 24 h after surgery as determined by flow cytometry. (D) Total Ly6G^+^ neutrophils, (D) total F4/80^+^ macrophages, and (E) total CD3^+^ T cells in the bronchoalveolar lavage fluids (BALF) as determined by flow cytometry 24 h after surgery. Results are represented as mean ± SEM of at least 5 mice per group. *p < 0.05, *** p < 0.001 and **** p < 0.0001.

Moreover, surface protein expression of ICAM-1 on endothelial cells was also significantly reduced after intranasal apratastat administration (Fig. 7). Together these data show that intranasal administration (e.g. inhalation) is also an efficient route of application of apratastat to combat Covid-19-associated lung inflammation.

**Figure 7.**
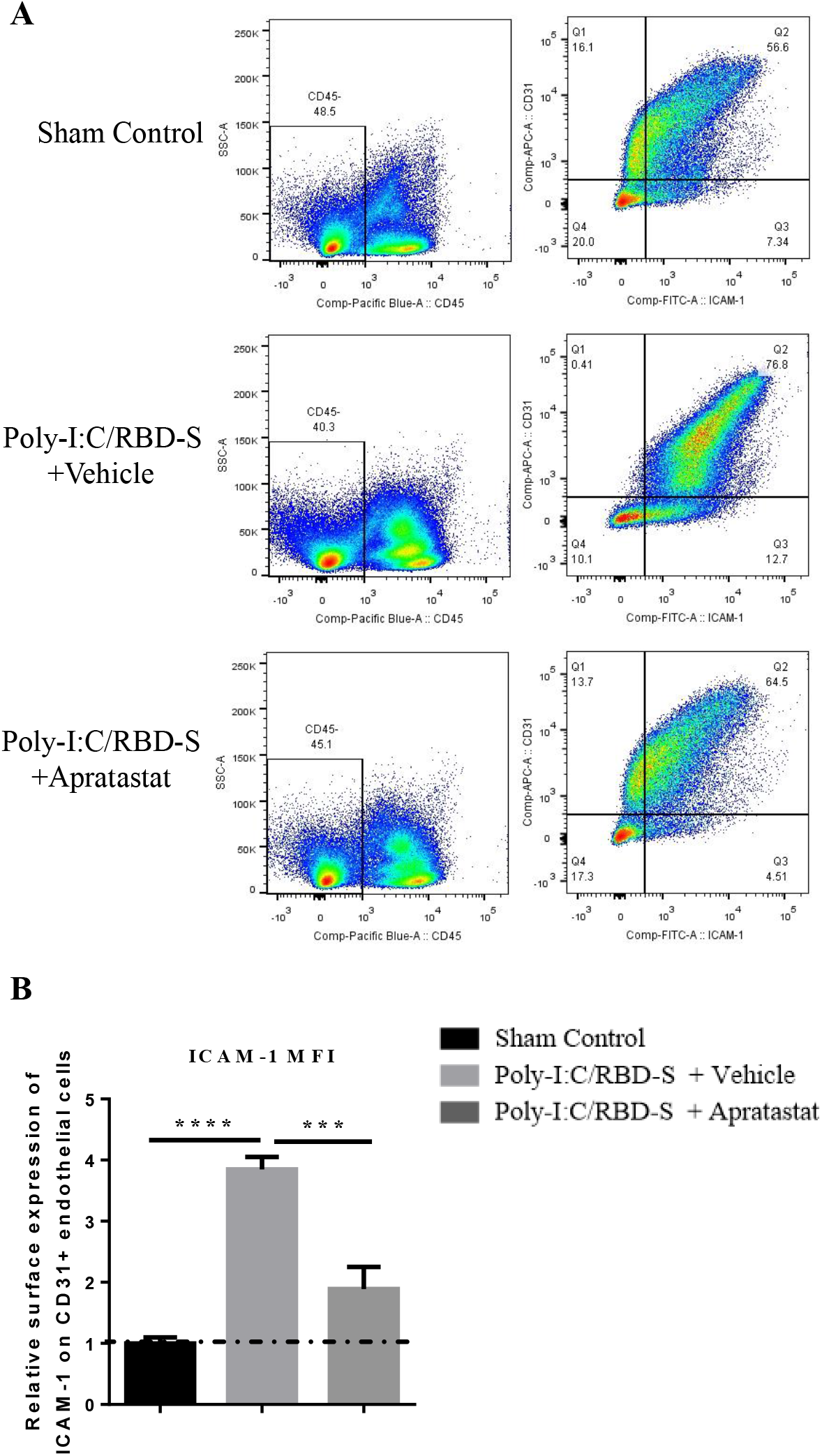
ICAM-1 expression in lung endothelial cells is reduced after intranasal administration of apratastat. (A) Representative ICAM1/CD31 plots from lung show a clear shift towards higher ICAM-1 levels in CD31 positive endothelial cells in poly-I:C/RBD-S-inflamed and vehicle-treated lungs. This shift is reverted in apratastat-treated inflamed mice. (B) Quantification of the MFI of the ICAM-1 signal from CD31^+^ICAM1^+^CD45^-^ cells Results are represented as mean ± SEM of at least 5 mice per group. i.n: intranasal. *** p < 0.001 and **** p < 0.0001.

## Discussion

Albeit herculean research efforts worldwide to understand Covid-19, it is still not entirely clear under which circumstances a SARS-CoV-2-infected individual develops severe Covid-19. Risk factors include age, comorbidities such as diabetes and obesity, and smoking, but sometimes even healthy young adults develop severe Covid-19 and die. Thus, specific and efficient treatments are desperately needed. One obstacle for research is that working with the virus itself requires high biosafety level infrastructure that is not available at many research facilities. Another obstacle is that mice, the most common animal in pre-clinical research, is not very susceptible to SARS-CoV-2 infection due to sequence differences (His 353) causing a lower affinity of mouse ACE2 for the spike protein (31). Thus, mouse models that can be used to study the molecular mechanisms related to Covid-19 without using the actual virus have been developed (19, 31-33). Here, we used a simple mouse model based on the use of the TLR3 ligand Poly I:C (19, 22) and the recombinant RBD of the SARS-CoV-2 spike protein in C57Bl/6 mice. We showed that these mice developed severe but not lethal lung injury with features characteristic for Covid-19-associated lung injury such as edema, fibrosis, and leukocyte infiltration. Importantly, these mice also had significantly increased NLR, another feature of severe Covid-19. These findings show that this simplified mouse model is convenient to study molecular mechanisms of Covid-19 pathogenesis. Here, we only used relatively young mice (8-10 weeks). It will be interesting to see whether older or obese animals will develop the more severe disease because if so, this model will also be appropriate to study the molecular differences in high-risk groups.

ACE2 is not just the receptor for SARS-CoV and SARS-CoV-2 (7, 34), but it is also a critical enzyme regulating the production of angiotensinogenic peptides, inflammation, and vascular and epithelial permeability (8). Thus, correct regulation of its enzymatic functions is of utmost importance for correct lung functionality, and loss of ACE2 by enzymatic cleavage is detrimental for lung functions (29). Two proteases have been shown to cleave ACE2, i.e. TMPRSS2 and ADAM17 (7, 12, 34). While ACE2 cleavage by TMPRSS2 is critical for SARS-CoV-2 entry into lung epithelial cells, the relevance of ADAM17-mediated ACE2 shedding is less clear. Here, we show that ADAM17 has a critical role in mediating Covid-19-related lung inflammation. Our data clearly show that ADAM17 inhibition ameliorates lung inflammation by attenuating the cytokine storm and neutrophil infiltration into the lung. ADAM17 is a well-known metalloproteinase expressed on the surface of many cells including lung epithelial cells and neutrophils that gets activated in response to many stimuli including coronavirus infections (35). ADAM17, also known as TNF-α-converting enzyme (TACE), is responsible for producing the active form of TNF-α (11). We showed that TNF-α production is indeed significantly reduced after ADAM17 inhibition in our Covid-19 model. Reduced TNF-α levels in turn may lead to reduced synthesis of other proinflammatory cytokines due to less NF-κB activation. This is likely an important mechanism responsible for the protective effect of ADAM17 inhibition against Covid-19-associated lung injury. Whether ADAM17 directly affects the production of other pro-inflammatory cytokines and chemokines needs to be investigated in the future. and thus contributes to the cytokine storm in severe Covid-19. Moreover, ADAM17 is also expressed in neutrophils, where it gets activated in response to integrin activation leading to cleavage of L-selectin, enhanced neutrophil effector functions and bacterial clearance in the lung (36). It will be important to analyze whether such mechanisms, at least to some degree, also play a role in viral clearance in mild to moderate Covid-19. However, in the context of severe Covid-19 with it is rather important to dampen neutrophil hyperactivation in response to the cytokine storm because excessive uncontrolled neutrophil effector functions including release of proteases, reactive oxygen species and NETs contribute to tissue damage as those substances do not distinguish between host and pathogens (5). On the other hand, lack of ADAM17 in neutrophils (37, 38) or ADAM17 inhibition in the context of sepsis (15) has been shown to accelerate neutrophil recruitment. We observed here significantly reduced neutrophil presence in lung tissue and BALF and consequently an improved NLR, but we cannot explain yet whether this inhibitory effect is due to ADAM17 inhibition in neutrophils or endothelial cells or both. It is also important to consider that neutrophil recruitment into the lung follows molecular steps different from those in other tissues (39) and that recruitment to different lung compartments in different Covid-19 stages is also governed by distinct mechanisms (40). Thus, more research is required to unravel the mechanisms underlying the reduced neutrophil recruitment patterns in our model of Covid-19-related lung inflammation.

Apratastat (TMI-005) and TMI-1 are structurally related orally bioavailable ADAM17 inhibitors of the thiomorpholine sulfonamide hydroxymate family (21). These compounds are highly specific for ADAM17 and barely inhibit the related ADAM10 (41). However, they also inhibit other matrix-metalloproteases (MMP) to a certain extent. Thus, it cannot be excluded that some of the observed effects also results from partial MMP inhibition. These inhibitors have been shown to inhibit TNF-α secretion in pre-clinical models of rheumatoid arthritis (RA) and have therefore been tested in clinical trials to treat RA patients (42). However, although effective in reducing TNF-α levels, these drugs did not improve the clinical manifestations in RA patients so that the study was terminated (43). Despite this disappointing clinical result, apratastat and TMI-1 could still be useful for the treatment of Covid-19 patients because in this context we observed not only a protective effect on TNF-α production, but also on neutrophil recruitment and NLR, all critical features of severe Covid-19 pathogenesis. Moreover, pharmacodynamics and pharmacokinetics data from clinical studies showed that at least apratastat is well tolerated and bioavailable in humans (44), so that it will be feasible to test this drug in clinical trials for efficacy in inhibiting severe Covid-19 development in SRAS-CoV-2-infected patients.

In summary, we provide experimental evidence that ADAM17 inhibition by apratastat and TMI-1 protects against lung inflammation in a novel, simplified mouse model of Covid-19-associated lung injury. ADAM17 reduced the cytokine storm and excessive neutrophil recruitment to the lung while improving NLR. Thus, we propose to use these drugs in clinical trials to test their efficacy in Covid-19 patients.

## Acknowledgement

This work was supported by funds from the Agencia Mexicana de Cooperación Internacional para el Desarrollo (AMEXCID) of the Secretaría de Relaciones Exteriores (SRE) (Projects AMEXCID_2020-4 to MS and AMEXCID_2020-3 to EMR)

MS acknowledges funding from Consejo Nacional de Ciencia y Tecnología (CONACYT; CB-2016-284292; and PRONAII Leukemia grant 302978 of the National Strategic Health Program). EMR acknowledges funding from CONACYT (CB2017-2018: A1-S-10743; Apoyo para proyectos de investigación científica, desarrollo tecnológico e innovación en salud ante la contingencia por covid-19: 311835), SEP-CINVESTAV (2018:1), and Gobierno del Estado de Hidalgo (Proyectos Sincrotrón 20201120 and 20201039).

SVR received a postdoctoral fellowship from CONACYT. NLL, KEJC, IMGF, RMC, AMG, JGFV, DRV and DIZV received a pre-doctoral fellowship from CONACYT.

## References

1. Zhu, F. C., X. H. Guan, Y. H. Li, J. Y. Huang, T. Jiang, L. H. Hou, J. X. Li, B. F. Yang, L. Wang, W. J. Wang, S. P. Wu, Z. Wang, X. H. Wu, J. J. Xu, Z. Zhang, S. Y. Jia, B. S. Wang, Y. Hu, J. J. Liu, J. Zhang, X. A. Qian, Q. Li, H. X. Pan, H. D. Jiang, P. Deng, J. B. Gou, X. W. Wang, X. H. Wang, and W. Chen. 2020. Immunogenicity and safety of a recombinant adenovirus type-5-vectored COVID-19 vaccine in healthy adults aged 18 years or older: a randomised, double-blind, placebo-controlled, phase 2 trial. Lancet 396: 479–488.

2. Chen, N., M. Zhou, X. Dong, J. Qu, F. Gong, Y. Han, Y. Qiu, J. Wang, Y. Liu, Y. Wei, J. Xia, T. Yu, X. Zhang, and L. Zhang. 2020. Epidemiological and clinical characteristics of 99 cases of 2019 novel coronavirus pneumonia in Wuhan, China: a descriptive study. Lancet 395: 507–513.

3. Tatum, D., S. Taghavi, A. Houghton, J. Stover, E. Toraih, and J. Duchesne. 2020. Neutrophil-to-Lymphocyte Ratio and Outcomes in Louisiana COVID-19 Patients. Shock 54: 652–658.

4. Vadillo, E., K. Taniguchi-Ponciano, C. Lopez-Macias, R. Carvente-Garcia, H. Mayani, E. Ferat-Osorio, G. Flores-Padilla, J. Torres, C. R. Gonzalez-Bonilla, A. Majluf, A. Albarran-Sanchez, J. C. Galan, E. Pena-Martinez, G. Silva-Roman, S. Vela-Patino, A. Ferreira-Hermosillo, C. Ramirez-Renteria, N. A. Espinoza-Sanchez, R. Pelayo-Camacho, L. Bonifaz, L. Arriaga-Pizano, C. Mata-Lozano, S. Andonegui-Elguera, N. Wacher, F. Blanco-Favela, R. De-Lira-Barraza, H. Villanueva-Compean, A. Esquivel-Pineda, R. Ramirez-Montes-de-Oca, C. Anda-Garay, M. Noyola-Garcia, L. Guizar-Garcia, A. Cerbulo-Vazquez, H. Zamudio-Meza, D. Marrero-Rodriguez, and M. Mercado. 2020. A Shift Towards an Immature Myeloid Profile in Peripheral Blood of Critically Ill COVID-19 Patients. Arch Med Res.

5. Narasaraju, T., B. M. Tang, M. Herrmann, S. Muller, V. T. K. Chow, and M. Radic. 2020. Neutrophilia and NETopathy as Key Pathologic Drivers of Progressive Lung Impairment in Patients With COVID-19. Front Pharmacol 11: 870.

6. Matthay, M. A., L. B. Ware, and G. A. Zimmerman. 2012. The acute respiratory distress syndrome. J Clin Invest 122: 2731–2740.

7. Hoffmann, M., H. Kleine-Weber, S. Schroeder, N. Kruger, T. Herrler, S. Erichsen, T. S. Schiergens, G. Herrler, N. H. Wu, A. Nitsche, M. A. Muller, C. Drosten, and S. Pohlmann. 2020. SARS-CoV-2 Cell Entry Depends on ACE2 and TMPRSS2 and Is Blocked by a Clinically Proven Protease Inhibitor. Cell 181: 271–280 e278.

8. Gheblawi, M., K. Wang, A. Viveiros, Q. Nguyen, J. C. Zhong, A. J. Turner, M. K. Raizada, M. B. Grant, and G. Y. Oudit. 2020. Angiotensin-Converting Enzyme 2: SARS-CoV-2 Receptor and Regulator of the Renin-Angiotensin System: Celebrating the 20th Anniversary of the Discovery of ACE2. Circ Res 126: 1456–1474.

9. Cao, Y., Y. Liu, J. Shang, Z. Yuan, F. Ping, S. Yao, Y. Guo, and Y. Li. 2019. Ang-(1-7) treatment attenuates lipopolysaccharide-induced early pulmonary fibrosis. Lab Invest 99: 1770–1783.

10. Kuba, K., Y. Imai, S. Rao, H. Gao, F. Guo, B. Guan, Y. Huan, P. Yang, Y. Zhang, W. Deng, L. Bao, B. Zhang, G. Liu, Z. Wang, M. Chappell, Y. Liu, D. Zheng, A. Leibbrandt, T. Wada, A. S. Slutsky, D. Liu, C. Qin, C. Jiang, and J. M. Penninger. 2005. A crucial role of angiotensin converting enzyme 2 (ACE2) in SARS coronavirus-induced lung injury. Nat Med 11: 875–879.

11. Zunke, F., and S. Rose-John. 2017. The shedding protease ADAM17: Physiology and pathophysiology. Biochim Biophys Acta Mol Cell Res 1864: 2059–2070.

12. Patel, V. B., N. Clarke, Z. Wang, D. Fan, N. Parajuli, R. Basu, B. Putko, Z. Kassiri, A. J. Turner, and G. Y. Oudit. 2014. Angiotensin II induced proteolytic cleavage of myocardial ACE2 is mediated by TACE/ADAM-17: a positive feedback mechanism in the RAS. J Mol Cell Cardiol 66: 167–176.

13. Cui, S. N., H. Y. Tan, and G. C. Fan. 2021. Immunopathological Roles of Neutrophils in Virus Infection and COVID-19. Shock.

14. Borges, L., T. C. Pithon-Curi, R. Curi, and E. Hatanaka. 2020. COVID-19 and Neutrophils: The Relationship between Hyperinflammation and Neutrophil Extracellular Traps. Mediators Inflamm 2020: 8829674.

15. Mishra, H. K., J. Ma, and B. Walcheck. 2017. Ectodomain Shedding by ADAM17: Its Role in Neutrophil Recruitment and the Impairment of This Process during Sepsis. Front Cell Infect Microbiol 7: 138.

16. Palau, V., M. Riera, and M. J. Soler. 2020. ADAM17 inhibition may exert a protective effect on COVID-19. Nephrol Dial Transplant 35: 1071–1072.

17. Kupferschmidt, K., and J. Cohen. 2020. Race to find COVID-19 treatments accelerates. Science 367: 1412–1413.

18. de la Cruz, J.J., L. Villanueva-Lizama, V. Dzul-Huchim, M. J. Ramirez-Sierra, P. Martinez-Vega, M. Rosado-Vallado, J. Ortega-Lopez, C. I. Flores-Pucheta, P. Gillespie, B. Zhan, M. E. Bottazzi, P. J. Hotez, and E. Dumonteil. 2019. Production of recombinant TSA-1 and evaluation of its potential for the immuno-therapeutic control of Trypanosoma cruzi infection in mice. Hum Vaccin Immunother 15: 210–219.

19. Gu, T., S. Zhao, G. Jin, M. Song, Y. Zhi, R. Zhao, F. Ma, Y. Zheng, K. Wang, H. Liu, M. Xin, W. Han, X. Li, C. D. Dong, K. Liu, and Z. Dong. 2020. Cytokine Signature Induced by SARS-CoV-2 Spike Protein in a Mouse Model. Front Immunol 11: 621441.

20. Ford, B. M., A. A. Eid, M. Gooz, J. L. Barnes, Y. C. Gorin, and H. E. Abboud. 2013. ADAM17 mediates Nox4 expression and NADPH oxidase activity in the kidney cortex of OVE26 mice. Am J Physiol Renal Physiol 305: F323–332.

21. Zhang, Y., J. Xu, J. Levin, M. Hegen, G. Li, H. Robertshaw, F. Brennan, T. Cummons, D. Clarke, N. Vansell, C. Nickerson-Nutter, D. Barone, K. Mohler, R. Black, J. Skotnicki, J. Gibbons, M. Feldmann, P. Frost, G. Larsen, and L. L. Lin. 2004. Identification and characterization of 4-[[4-(2-butynyloxy)phenyl]sulfonyl]-N-hydroxy-2,2-dimethyl-(3S)thiomorpholinecar boxamide (TMI-1), a novel dual tumor necrosis factor-alpha-converting enzyme/matrix metalloprotease inhibitor for the treatment of rheumatoid arthritis. J Pharmacol Exp Ther 309: 348–355.

22. Gan, T., Y. Yang, F. Hu, X. Chen, J. Zhou, Y. Li, Y. Xu, H. Wang, Y. Chen, and M. Zhang. 2018. TLR3 Regulated Poly I:C-Induced Neutrophil Extracellular Traps and Acute Lung Injury Partly Through p38 MAP Kinase. Front Microbiol 9: 3174.

23. Sun, F., G. Xiao, and Z. Qu. 2017. Murine Bronchoalveolar Lavage. Bio Protoc 7.

24. Klopfleisch, R. 2013. Multiparametric and semiquantitative scoring systems for the evaluation of mouse model histopathology--a systematic review. BMC Vet Res 9: 123.

25. Schaller, T., K. Hirschbuhl, K. Burkhardt, G. Braun, M. Trepel, B. Markl, and R. Claus. 2020. Postmortem Examination of Patients With COVID-19. JAMA 323: 2518–2520.

26. Li, X., C. Liu, Z. Mao, M. Xiao, L. Wang, S. Qi, and F. Zhou. 2020. Predictive values of neutrophil-to-lymphocyte ratio on disease severity and mortality in COVID-19 patients: a systematic review and meta-analysis. Crit Care 24: 647.

27. Hussman, J. P. 2020. Cellular and Molecular Pathways of COVID-19 and Potential Points of Therapeutic Intervention. Front Pharmacol 11: 1169.

28. Xiong, Y., Y. Liu, L. Cao, D. Wang, M. Guo, A. Jiang, D. Guo, W. Hu, J. Yang, Z. Tang, H. Wu, Y. Lin, M. Zhang, Q. Zhang, M. Shi, Y. Liu, Y. Zhou, K. Lan, and Y. Chen. 2020. Transcriptomic characteristics of bronchoalveolar lavage fluid and peripheral blood mononuclear cells in COVID-19 patients. Emerg Microbes Infect 9: 761–770.

29. Zipeto, D., J. D. F. Palmeira, G. A. Arganaraz, and E. R. Arganaraz. 2020. ACE2/ADAM17/TMPRSS2 Interplay May Be the Main Risk Factor for COVID-19. Front Immunol 11: 576745.

30. Vestweber, D. 2015. How leukocytes cross the vascular endothelium. Nat Rev Immunol 15: 692–704.

31. Ren, W., Y. Zhu, Y. Wang, H. Shi, Y. Yu, G. Hu, F. Feng, X. Zhao, J. Lan, J. Wu, D. J. Kenney, F. Douam, Y. Tong, J. Zhong, Y. Xie, X. Wang, Z. Yuan, D. Zhou, R. Zhang, and Q. Ding. 2021. Comparative analysis reveals the species-specific genetic determinants of ACE2 required for SARS-CoV-2 entry. PLoS Pathog 17: e1009392.

32. Dinnon, K. H., 3rd, S. R. Leist, A. Schafer, C. E. Edwards, D. R. Martinez, S. A. Montgomery, A. West, B. L. Yount, Jr., Y. J. Hou, L. E. Adams, K. L. Gully, A. J. Brown, E. Huang, M. D. Bryant, I. C. Choong, J. S. Glenn, L. E. Gralinski, T. P. Sheahan, and R. S. Baric. 2020. A mouse-adapted model of SARS-CoV-2 to test COVID-19 countermeasures. Nature 586: 560–566.

33. McCray, P. B., Jr., L. Pewe, C. Wohlford-Lenane, M. Hickey, L. Manzel, L. Shi, J. Netland, H. P. Jia, C. Halabi, C. D. Sigmund, D. K. Meyerholz, P. Kirby, D. C. Look, and S. Perlman. 2007. Lethal infection of K18-hACE2 mice infected with severe acute respiratory syndrome coronavirus. J Virol 81: 813–821.

34. Heurich, A., H. Hofmann-Winkler, S. Gierer, T. Liepold, O. Jahn, and S. Pohlmann. 2014. TMPRSS2 and ADAM17 cleave ACE2 differentially and only proteolysis by TMPRSS2 augments entry driven by the severe acute respiratory syndrome coronavirus spike protein. J Virol 88: 1293–1307.

35. Haga, S., N. Yamamoto, C. Nakai-Murakami, Y. Osawa, K. Tokunaga, T. Sata, N. Yamamoto, T. Sasazuki, and Y. Ishizaka. 2008. Modulation of TNF-alpha-converting enzyme by the spike protein of SARS-CoV and ACE2 induces TNF-alpha production and facilitates viral entry. Proc Natl Acad Sci U S A 105: 7809–7814.

36. Cappenberg, A., A. Margraf, K. Thomas, B. Bardel, D. A. McCreedy, V. Van Marck, A. Mellmann, C. A. Lowell, and A. Zarbock. 2019. L-selectin shedding affects bacterial clearance in the lung: a new regulatory pathway for integrin outside-in signaling. Blood 134: 1445–1457.

37. Long, C., Y. Wang, A. H. Herrera, K. Horiuchi, and B. Walcheck. 2010. In vivo role of leukocyte ADAM17 in the inflammatory and host responses during E. coli-mediated peritonitis. J Leukoc Biol 87: 1097–1101.

38. Tang, J., A. Zarbock, I. Gomez, C. L. Wilson, C. T. Lefort, A. Stadtmann, B. Bell, L. C. Huang, K. Ley, and E. W. Raines. 2011. Adam17-dependent shedding limits early neutrophil influx but does not alter early monocyte recruitment to inflammatory sites. Blood 118: 786–794.

39. Margraf, A., K. Ley, and A. Zarbock. 2019. Neutrophil Recruitment: From Model Systems to Tissue-Specific Patterns. Trends Immunol 40: 613–634.

40. Alon, R., M. Sportiello, S. Kozlovski, A. Kumar, E. C. Reilly, A. Zarbock, N. Garbi, and D. J. Topham. 2021. Leukocyte trafficking to the lungs and beyond: lessons from influenza for COVID-19. Nat Rev Immunol 21: 49–64.

41. Ludwig, A., C. Hundhausen, M. H. Lambert, N. Broadway, R. C. Andrews, D. M. Bickett, M. A. Leesnitzer, and J. D. Becherer. 2005. Metalloproteinase inhibitors for the disintegrin-like metalloproteinases ADAM10 and ADAM17 that differentially block constitutive and phorbol ester-inducible shedding of cell surface molecules. Comb Chem High Throughput Screen 8: 161–171.

42. Moss, M. L., L. Sklair-Tavron, and R. Nudelman. 2008. Drug insight: tumor necrosis factor-converting enzyme as a pharmaceutical target for rheumatoid arthritis. Nat Clin Pract Rheumatol 4: 300–309.

43. Thabet, M. M., and T. W. Huizinga. 2006. Drug evaluation: apratastat, a novel TACE/MMP inhibitor for rheumatoid arthritis. Curr Opin Investig Drugs 7: 1014–1019.

44. Shu, C., H. Zhou, M. Afsharvand, L. Duan, H. Zhang, R. Noveck, and D. Raible. 2011. Pharmacokinetic-pharmacodynamic modeling of apratastat: a population-based approach. J Clin Pharmacol 51: 472–481.

